# PEZO-1 and TRP-4 mechanosensors are involved in mating behavior in *Caenorhabditis elegans*

**DOI:** 10.1101/2022.08.31.506045

**Authors:** Katherine I. Brugman, Vladislav Susoy, Allyson J. Whittaker, Wilber Palma, Stephanie Nava, Aravinthan D.T. Samuel, Paul W. Sternberg

**Author notes:** correspondence (PWS). equal contribution (Katherine I. Brugman, Vladislav Susoy).

## Abstract

Male mating in *Caenorhabditis elegans* is a complex behavior with a strong mechanosensory component. *C. elegans* has several characterized mechanotransducer proteins, but few have been shown to contribute to mating. Here, we investigated the roles of PEZO-1, a piezo channel, and TRP-4, a mechanotransducing TRPN channel, in male mating behavior. We show that *pezo-1* is expressed in several male-specific neurons with known roles in mating. We show that, among other neurons, *trp-4* is expressed in the PCA sensory neuron, which monitors relative sliding between the male and the hermaphrodite and inhibits neurons involved in vulva detection. Mutations in both genes compromise many steps of mating, including initial response to the hermaphrodite, scanning, turning, and vulva detection. We performed pan-neuronal imaging during mating between freely-moving mutant males and hermaphrodites. Both *pezo-1* and *trp-4* mutants showed spurious activation of the sensory neurons involved in vulva detection. In *trp-4* mutants, this spurious activation might be caused by PCA failure to inhibit vulva-detecting neurons during scanning. Indeed, we show that without functional TRP-4, PCA fails to detect the relative sliding between the male and hermaphrodite. Cell-specific TRP-4 expression restores PCA’s mechanosensory function. Our results demonstrate new roles for both PEZO-1 and TRP-4 mechanotransducers in *C. elegans* mating behavior.

## INTRODUCTION

Mechanosensory transduction converts diverse mechanical stimuli into biological responses from the workings of cochlear hair cells to osmotic pressure regulators in bacteria to proprioception and touch ^1–4^. Mechanosensation is adapted to rapidly changing stimuli through the direct sensitivity of diverse ionotropic receptors^5^. Several primary mechanotransducers have been identified in *C. elegans* including a TRPA ortholog TRPA-1, a TRPV ortholog OSM-9, a TRPN ortholog TRP-4, and a DeG/ENaC channel composed of MEC-4/MEC-10 and accessory proteins. These channels regulate diverse rapid behavioral responses ^6–8^.

*C. elegans* also expresses PEZO-1, an invertebrate homologue of the piezo class of mechanosensory proteins. Piezo proteins comprise non-selective and conserved cation channels found in plants, animals, and fungi^9–14^. All piezo proteins form a homotrimeric structure with a cation-selective central pore surrounded by three N-terminal propeller blades and a cap that covers the pore^15–17^. Piezo proteins have diverse mechanosensory roles from nociception in *Drosophila* to blood cell volume regulation in mice, zebrafish, and humans^10,13,18–25^. Like other invertebrates, *C. elegans* has a single piezo protein, PEZO-1. In the hermaphrodite, PEZO-1 has been shown to have roles in ovulation, defecation, and food intake^11,26–28^.

Ionotropic channels in the TRP superfamily are composed of six transmembrane domain proteins that form large cation-selective pore-forming tetramers^4,29^. TRP channels are divided into six subfamilies based on function and sequence identity^4^. TRP channels are implicated in diverse sensory modalities including thermosensation, mechanosensation, taste, smell, and hearing^4,30–35^. The *C. elegans* TRP-4 channel is part of the mechanosensative TRPN subfamily, and has been implicated in locomotion: *trp-4* mutant animals exhibit exaggerated body bending, likely due to a defect in proprioception mediated by the TRP-4 expressing DVA neuron^7,30^.

*C. elegans* is a self-fertilizing species with occasional males^36–39^. Mating in the *C. elegans* male is a complex behavior, which involves several different steps, or behavioral motifs, including initial attraction, scanning for the vulva, turning, stopping at the vulva, insertion of the spicules and release of sperm^40,41^. Each step exhibits distinct dynamics. Scanning starts when the male contacts the hermaphrodite with the ventral part of his tail and backs along her cuticle. Turning occurs whenever the male reaches the hermaphrodite head or tail during scanning, and allows him to continue scanning on her other side. The male stops when he reaches the vulva. Vulva recognition is followed by prodding when he rapidly presses his spicules (sclerotized structures for mating) into her vulval region, along with fine-scale location of the vulva opening^42^. Successful spicule insertion triggers sperm release. The dynamics of mating behavior is driven by diverse patterns of sensory recognition of the hermaphrodite by neurons in the male tail^40,43^.

Many male sensory neurons appear to be polymodal with both chemosensory and mechanosensory functions. For example, the hook neurons HOA and HOB have ciliated sensory endings but are also embedded in a structure with a mechanosensory morphology (the hook)^44^. The ciliated ray neurons (R[1-9]A and R[1-9]B) are open to the exterior of the worm (except the R6 neurons), and are embedded in elongated sensory rays. The post-cloacal sensilla are a pair of sensory structures just posterior to the cloaca, and each contain the ciliated PCA, PCB, and PCC sensory neurons^44,45^. Laser ablation of these sensory structures results in a defect in particular motifs of mating^40^. Ablation of rays along with other ventral organs results in a male that fails to initiate scanning in response to ventral contact^40^. Ablation of either the hook sensillum or either associated neuron (HOA and HOB) results in a male that is unable to stop at the vulva normally, instead backing slowly with spicule partially extruded until contacting the vulva^40^. Ablation of postcloacal sensilla neurons results in a male with nearly wild-type mating ability except that he tends to ‘lose’ the vulva more often^40^. Ablation of both the hook sensillum and post cloacal sensillum results in a male that is unable to stop at the vulva or locate the vulva through slow searching^40^.

A prior study confirmed the role of one of the known mechanoreceptors, the MEC-4/MEC-10 complex (DeG/ENaC channel) in mating. Specifically, *mec-4* gain-of-function, *mec-4* reduction-of-function, and *mec-10* loss-of-function mutants have turning defects, suggesting the involvement of the MEC-4/MEC-10 complex in regulating turning behavior during mating^8^. Another study demonstrated that PKD-1 and PKD-2, *C. elegans* homologues of human PKD1 and PKD2, are involved in mating behaviors with mechanosensory components – initial response to the hermaphrodite and vulva detection^46^. PKD1 is thought to have mechanosensory properties^47^. Here, we examine two other potential mechanoreceptors, namely the piezo mechanoreceptor in *C. elegans*, PEZO-1, and the transient receptor potential channel protein, TRP-4, which is expressed in PCA and several ray neurons. We generate mutants defective in *pezo-1* and *trp-4*, and establish their contribution to male mating by analyzing behavior and neuronal activity in freely-moving males.

## METHODS

### General methods

*C. elegans* strains were cultured at 23°C^41^. Assays were also conducted at this temperature, except the egg-laying assay which was performed at 20°C.

### *C. elegans* strains

This study

PS7489 *pezo-1(sy1113)* IV

PS8908 *pha-1* III; *him-5* V; *syEx1771* [pBX-1 + *pezo-1(p3)*::GFP]

PS8909 *pha-1* III; *him-5* V; *syEx1792* [pBX-1 + *pezo-1(p5)*::GFP]

PS8040 *hpIs675* [P*rgef*-1 GCaMP6::3xNLS::mNeptune + *lin-15*(+)]; *him-5 (e1490)* V

PS8041 *hpIs675* [P*rgef*-1 GCaMP6::3xNLS::mNeptune + *lin-15*(+)]; *pezo-1 (sy1113)* IV; *him-5 (e1490)* V

PS9157 *hpIs675* [P*rgef*-1 GCaMP6::3xNLS::mNeptune + *lin-15*(+)]; *pezo-1 (av240)* IV; *him-5 (e1490)* V

PS9045 *hpIs675* [P*rgef*-1 GCaMP6::3xNLS::mNeptune + *lin-15*(+)]; *trp-4 (sy695)* I; *him-5 (e1490)* V

PS9669 *syEx1715*[*(pWP082)Peat-4(p8)*::TRP-4::*unc-54* 3’UTR + unc-122::RFP]; *hpIs675*

[*Prgef-1* GCaMP6::3xNLS::mNeptune + *lin-15*(+)]; *trp-4 (sy695)* I; *him-5 (e1490)* V

Other strains

CB4088 *him-5 (e1490)* V

AG570 *pezo-1(av240)* IV

PS1905 *trp-4 (sy695)* I; *him-5 (e1490)* V

ZM9624 *hpIs675* [P*rgef*-1 GCaMP6::3xNLS::mNeptune] X

ADS1002 *aeaIs010* [P*rgef*-1::GCaMP6s::3xNLS + *lin-15*(+)]

BB92 *uuEx18* [*dcr-1*(wild-type) + *dpy-30*::mCherry]

ADS1014 *otIs377* [*myo-3*p::mCherry]; [*unc-64*(e246)](https://cgc.umn.edu/strain/search?st1=e246&sf1=all) III

OH10235 *bxIs19* [*trp-4*p::GFP]

OH13105 *him-5* (e1490) *otIs564* [*unc-47*(fosmid)::SL2::H2B::mChopti + *pha-1*(+)] V

OH13645 *pha-1* (e2123) III; *him-5*(e1490) V; *otIs518* [*eat-4*(fosmid)::SL2::mCherry::H2B + *pha-1*(+)]

OH10972 *otIs376* [*eat-4(prom4)*::GFP + *rol-6(su1006)*]

OH11152 *otIs392* [*eat-4(prom6)*::GFP + *ttx-3::DsRed*]

OH16719 *otIs138* [*ser-2(prom3)*::GFP *+rol-6(su1006)*] X; *otIs521* [*eat-4(prom8)*::tagRFP + *ttx-3*::GFP]

OH14018 *otIs520* [*eat-4(prom11)*::GFP + *ttx-3::mCherry*]

### *pezo-1::GFP* transcriptional fusions

To visualize the cellular expression patterns of *pezo-1*, a transcriptional fusion plasmid of several predicted regulatory regions and GFP were generated and injected into N2 hermaphrodites along with a *myo-2* DsRed co-injection marker (Figure 1A,B). Several suspected regulatory regions were selected, including the region 2 kb preceding the ATG site of the primary isoform as well as those of proposed alternate isoforms. A combined transcriptional fusion plasmid was generated from a 9 kb combination of the regions that exhibited the broadest expression patterns (TCCTACCTTGTGCCTGTCTACGTA…TCTGTTGTTGCTGTGTTACGACCA and GCACATGCTGCAGGTTAGTTTATT…AGGGTATCGCCTGAAAAAACAATG) in both *him-5* background with a *myo-2* DsRed co-injection marker and in *pha-1* background with a *pha-1* rescue plasmid as a marker. Expression patterns were examined with epifluorescence microscopy and a 100x objective.

**Figure 1.**
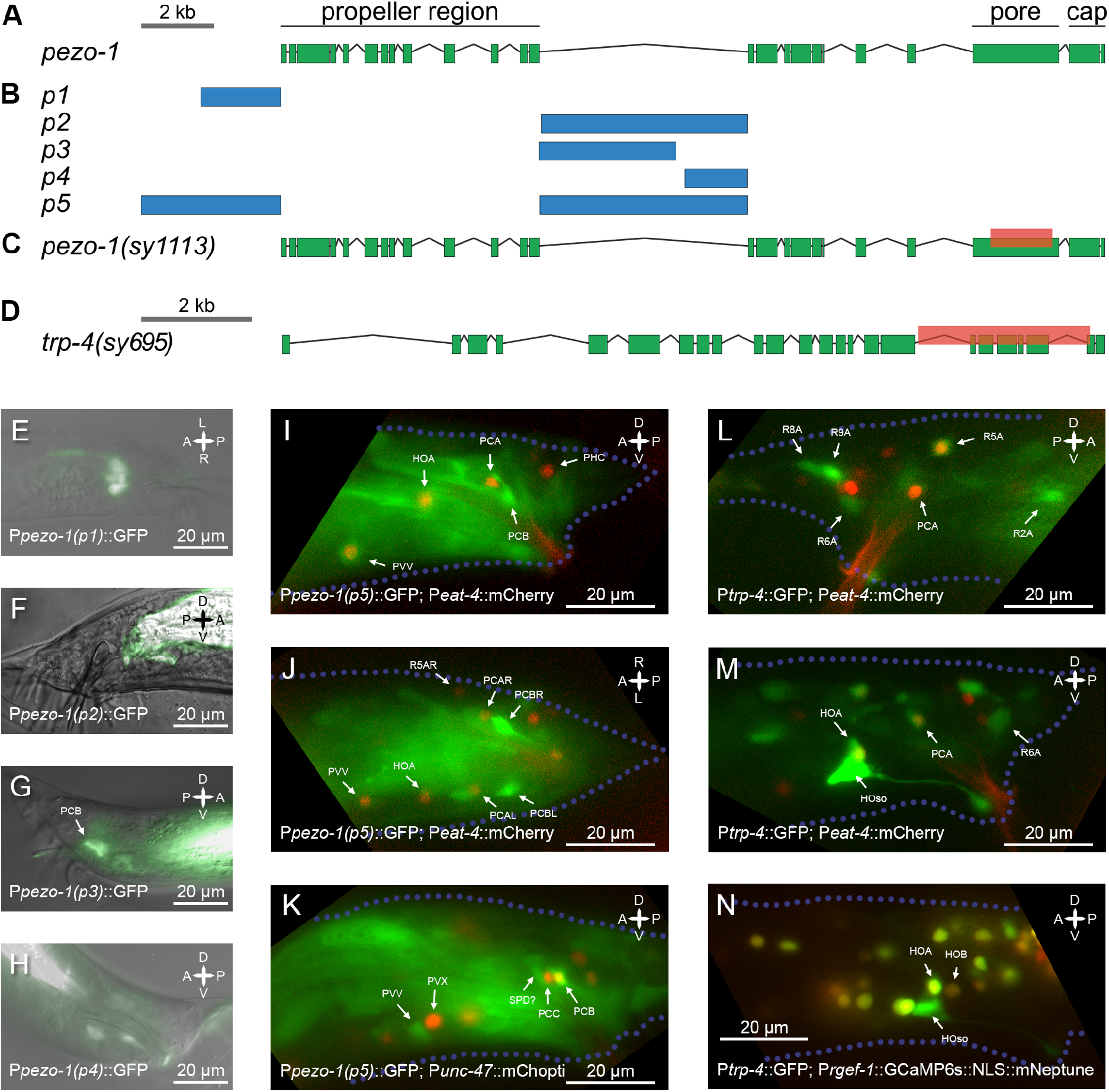
*pezo-1* and *trp-4* gene structure and expression. (A) Gene layout for *pezo-1* in *C. elegans*. (B) Non-coding regions used for creating transcriptional fusion reporters. *pezo-1(p5)* included two regulatory regions fused together. (C) A 2.1kb C-terminal deletion was generated using Crispr/Cas9. The deletion affected the pore domain of PEZO-1. (D) Gene layout for *trp-4*; mutant *trp-4* allele is shown. (E) GFP expression of transcriptional fusion using *pezo-1(p1)*, a region 2kb upstream of the start codon. Only some intestinal glands express GFP. (F) GFP expression under *pezo-1(p2)*, which includes most of the central intron minus ∼100 bp at the 5’ end. Only intestinal expression is visible. (G) GFP expression under *pezo-1(p3)*, which includes the front half of the central intron. Neuronal expression is visible, including expression in the PCB sensory neuron. (H) Expression pattern of *pezo-1(p4)*, which captures the back half of the central intron. Neuronal expression is visible. (I-K) GFP expression for *pezo-1(p5)*, which includes the fusion of two regulatory regions, is shown, coexpressed with P*eat-4::*mCherry (I and J) and P*unc-47::*mChopti (K). PCB and PVV, which show strong *pezo-1::*GFP expression, are shown in relation to the marker strains. (L - M) Glutamatergic PCA shows *trp-4::*GFP expression. PCA identity is confirmed based on the expression of *eat-4*, a PCA marker. Strong expression of *trp-4*::GFP is also observed in the hook socket cell and a weak *trp-4*::GFP expression can be seen in the HOA sensory neuron.

### CRISPR/Cas9 deletion mutant

To generate the 2kb deletion mutant, *pezo-1(sy1113)*, we injected a 46168 Crispr/Cas9 plasmid (*peft-3*::Cas9-sv40_NLS::*ttb-2* UTR) along with pRB1017 plasmids (pU6::gRNA construct) containing our guide RNA sequences (CAGAAGCTCGTAAGCCAGG, AGGTCGAGGTCGTGAGCGG, CCACCACTTTACGAGATGG, ATAGGCAGCTTCGAACTGG) into young adult hermaphrodites^48,49^. In addition, we co-injected a *dpy-10* guide RNA plasmid, also inserted into pRB1017, and a repair oligo with the *dpy-10* mutation as a marker which was later crossed out prior to use. Successful injections were identified by the presence of Dumpy or Roller progeny. 120 Dumpy and Roller progeny were singled onto individual plates and screened for the deletion mutation. Worm lysate was split between a control PCR with one internal and external primer (GGCGACCGTTGGTTGGGCTGGTTT, AATACGAGAGCCTTCACATCATC) and a detection PCR with two primers external to the proposed deletion (GGCGACCGTTGGTTGGGCTGGTTT, ATCCTGTGTCCGATCCTGACG). A homozygous mutation line was determined by the absence of a band in the control PCR and the presence of a 500 bp band in the detection PCR (Figure 1C). The strain was then outcrossed and crossed into *him-5* to remove the Dumpy and Roller marker mutations and verified by PCR with the above primers. The deletion was also sequenced and verified to be in frame and at the location, IV:9347253…9345244.

### Imaging Strains

For pan-neuronal imaging we used ZM9624, a strain designed to co-express calcium indicator GCaMP6s and red fluorescent protein mNeptune in all neuronal nuclei under the *rgef-1* promoter^43^. PS8041 was generated by crossing ZM9624 into a *pezo-1(sy1113); him-5(e1490)* mutant background, PS8040 was generated by crossing ZM9624 into *him-5(e1490)*, and PS9045 was generated by crossing ZM9624 into *trp-4(sy695); him-5(e1490)*. To track mating events, we used hermaphrodites that expressed red fluorescent markers in their cuticle or muscles. To encourage the richness of male behavior we used partners with normal motility or showing only mild unc phenotypes.

### Cell-specific rescue of *trp-4*

We sought to rescue *trp-4(sy695)* mutation (Figure 1D) by expressing TRP-4 in the PCA neuron. PCA is a glutamatergic neuron expressing *eat-4*^50,51^. *eat-4* promoter is known to drive gene expression in multiple neurons. Previously, it was shown that different parts of the *eat-4* promoters drive expression in different subsets of glutamatergic neurons in the hermaphrodite^52,53^. We screened four existing marker lines to identify *eat-4* promoter regions that drive specific gene expression in PCA – OH10972, OH11152, OH16719, and OH14018^52,53^. We found that one line – OH16719, which included a 289 bp region starting 5327 bp upstream of the *eat-4* start codon, showed strong tagRFP expression in PCA and HOA neurons in the tail and 3 neurons in the head (OLL, OLQ, RIA, based on Serrano-Saiz et al. (2013)^52^). We used the 289 bp promoter and the coding sequence of *trp-4* to generate a rescue construct. The construct was directly injected, along with a co-injection marker into PS9045 imaging strain, which carried *trp-4(sy695)* mutation. The resulting strain – PS9669 – was screened for animals showing expression of the co-injection marker, which were selected for pan-neuronal imaging.

### Recording of neuronal activity in freely moving males

To record activity of neurons in freely-moving males we used a custom-built spinning disc confocal microscopy setup^43,54^. Briefly, virgin L4 males were picked onto a separate plate with OP50 and were kept on the plate for 15 to 20 hours before imaging. For the imaging experiments, a single virgin male was placed onto a 10 cm NGM agar plate containing a small amount of OP50 for food, 10 μl of NGM buffer, and 3 to 5 fluorescently labeled hermaphrodites. The plate was covered with a No1 coverslip. The animals were able to navigate freely under the coverslip. The volumetric imaging was performed with a 40x 0.95 NA oil objective (Nikon Plan Apo Lambda series) controlled by a piezoelectric stage. We imaged 10 brain volumes per second, each volume consisting of 20 optical slices approximately 1.75 micrometers apart. The emitted light from the objective was split into green and red channels and imaged with two separate Andor Zyla 4.2 sCMOS cameras at 200 Hz. Each camera recorded a 256×512 pixel region with 0.45 μm pixel size. The posterior nervous system of the male was tracked continuously throughout the imaging session. This was achieved by adjusting the microscope stage position in x, y, and z. Imaging experiments lasted from 1.5 to 10 minutes and specifically aimed at capturing scanning, turning and vulva detection events.

### Extraction of neuronal activity signals

Neuronal activity traces were extracted from the raw data following image preprocessing and registration. Registered data were down-sampled to 5 volumes per second. Neuronal segmentation and tracking were performed using manual and semi-automated methods with the help of Fiji plugin MaMuT 0.27^55,56^. Fluorescent signals from green and red channels were extracted for 2.25×2.25×3.5 μm regions of interest centered on neuronal nuclei. Savitzky-Golay filtering with polynomial order of 1 and frame length 13 was used for noise reduction^57^. To minimize motion artifacts, a ratiometric approach was used for calculating neuronal activity traces^43^.

For *pezo-1(sy1113), pezo-1(av240)*, and *trp-4(sy695)* males, we extracted activities of eight neurons implicated in vulva detection and scanning^40,43,58^. These neurons included post-cloacal sensilla neurons PCB, PCC, PCA, the hook neurons HOA, HOB, the spicule neuron SPC, the ray neuron R2B, and the interneuron PVX. In addition, we extracted activities of ray neurons R2A, R4A, and R6A for *trp-4* males and of the PVV interneuron for *pezo-1(av240)* males. For *trp-4* rescue males (PS9669), we quantified PCA activity. For control males, we used previously generated recordings of ZM9624 males, which were performed using the same microscopy setup^43^. From the recordings of mating of mutant and control males, we also tabulated discrete behavioral motifs expressed by the male, which included ventral contact with the hermaphrodite, vulva contact, successful and failed turning attempts and switching between backward and forward sliding relative to the hermaphrodite.

To quantify neuronal responses to the onsets of discrete behavioral motifs, we extracted neuronal activity traces aligned to the onsets of those specific behavioral motifs for a window spanning 7 seconds before and 13 seconds after each motif onset. Dataset-averaged response curves were calculated for each neuron and motif and normalized to a minimum value of 0 and a maximum value of 1. We calculated cross-correlations between these normalized traces and binarized behavioral event traces, with the lag parameter set to 5 (corresponds to 1 second). A time lag with the absolute maximum correlation across all motif instances was identified. One-sample t-test was used to test if the correlation between the neuron’s traces and behavior at that time lag was significantly greater than zero.

### Mating with freely-moving hermaphrodites

L4 males were picked onto a separate plate overnight and all experiments were performed on the next day. Mating plates were prepared by placing 2 μl of concentrated *E. coli* OP50 solution onto a 6 cm NGM agar plate and allowed to dry. Two *him-5* hermaphrodites were placed onto the OP50 spot and allowed to acclimate for 5 minutes. A single male was added to the plate and the plate was imaged with a Nikon microscope equipped with a 10x objective and a PointGrey camera. The male was tracked continuously by adjusting the stage position with a stage controller. The experiment was recorded at 5 frames per second. The recording started when the male approached a hermaphrodite, and continued for 15 minutes or until sperm release, whichever earlier. Mutant and control males were imaged in parallel.

Collected recordings of mating were analyzed blindly. For each experiment, we tabulated timestamps for the following behavioral events: (i) ventral contact of the male tale with the hermaphrodite body, (ii) loss of ventral contact, (iii) contact with the vulva, (iv) end of vulva contact, (v) successful turns, (vi) failed turn attempts, (viii) sliding over to lateral sides of the hermaphrodite, (vii) beginnings and ends of pauses away from the vulva, (viii) copulation and sperm release. From these timestamps we calculated several measures of mating performance: (i) turning success rate, (ii) the number of ejaculation events per ventral contact duration, (iii) the number of ejaculation events per vulva contact duration, (iv) the ratio between spurious pausing duration and vulva contact duration, (v) the ratio between passing the vulva events and ventral contact duration, (vi) mean vulva contact, (vii) the number of ‘sliding over’ events per ventral contact duration, and (viii) the number of ventral contact loss events per ventral contact duration.

### Body size measurements

Mutations in *pezo-1* were previously shown to cause feeding-related phenotypes in *C. elegans*^27,28^. To test whether *trp-4* and *pezo-1* mutants showed signs of starvation, we measured their body size. L4 males were picked onto a separate plate and imaged on the next day. The males were mounted on a glass slide with an agar pad and immobilized using sodium azide. Imaging was done with a 4x objective. We measured the surface area of the sagittal section of each male using Fiji^56^, as a proxy for body size (Supplemental Figure 1).

## RESULTS

### *pezo-1* and *trp-4* are expressed in male-specific neurons

To determine *pezo-1* expression, we generated five transcriptional fusion constructs that included different predicted regulatory regions for the *pezo-1* gene (Figure 1A,B). These constructs showed different but overlapping expression patterns (Figure 1E-H). Even small differences in the promoter region used for fusion constructs resulted in substantial changes in the expression pattern. For promoters 2 and 3, a 100 bp difference at the 5’ end coincided with a substantial difference in expression among male neurons, suggesting that this region is important for determining *pezo-1* expression. A fusion of the region 4kb upstream of *pezo-1* and its largest intron (Figure 1B, *pezo-1(p5)*) led to an expression pattern that contained all sparser expression patterns with other transcriptional fusions.

We observed *pezo-1* expression in several head neurons, pharynx-intestinal valve, vulval muscle, spermatheca, CAN, and male tail neurons. Male-specific neurons that showed strong *pezo-1* expression included PCB, PVV, and several sensory ray neurons (Figure 1G-K, see also Millet et al. (2021)^27^). PCB was previously shown to be required for vulva recognition^40^. PVV initiates sharp ventral bending during turning^43^. Ray neurons are required for hermaphrodite recognition and scanning^40,59^.

*trp-4* was previously shown to be expressed in eight pairs of A-type ray neurons^59^. We examined *trp-4* expression using the OH10235 *bxIs19 [trp-4p::GFP]* transcriptional fusion and noted that *trp-4* is also expressed in the hook neuron HOA and in the PCA postcloacal sensilla neuron (Figure 1L-N). HOA and PCA expression was confirmed using otIs518, an *eat-4* marker^51,60^. HOA is required for vulva recognition^40,43^. PCA becomes active when the male slides backwards relative to the hermaphrodite, and thus, PCA has been proposed to have mechanosensory function^43^. PCA also becomes active shortly after spicule insertion^43,51^, and, together with other postcloacal sensilla neurons, it has been implicated in vulva recognition^40,61^.

### *pezo-1* and *trp-4* mutants have impaired mating

Given that both *pezo-1* and *trp-4* are expressed in neurons with key roles in mating behavior, we wanted to test if mutations in *pezo-1* and *trp-4* compromise male mating. We quantified the mating efficiency of two *pezo-1* mutants (*pezo-1(sy1113)* and *pezo-1(av240)*), a *trp-4 (sy695)* mutant, and control males (*him-5*). Virgin males were allowed to mate for 15 minutes with freely-moving hermaphrodites. Freely-moving hermaphrodites allow a more complete analysis of behavioral phenotypes as they present greater challenges to successful mating. For each male, we calculated the mating success index: the number of ejaculation events per courtship duration. This index is higher when the male succeeds in mating after a short period of scanning. The index is lower when the male succeeds in mating after a long period of scanning or fails to mate at all. *pezo-1(av240)* and *trp-4(sy695)* mutants, but not *pezo-1(sy1113)* were significantly less efficient at mating compared to control males (Wilcoxon test, *p<0*.*037, p<0*.*045, p<0*.*45* respectively) (Figure 2A). Only 10% of *pezo-1(av240)* and 20% of *trp-4(sy695)* mutant males were able to mate successfully, compared to 60% and 50% of control and *pezo-1(sy1113)* males, respectively. We conclude that both *pezo-1* and *trp-4* mutations compromise mating efficiency.

**Figure 2.**
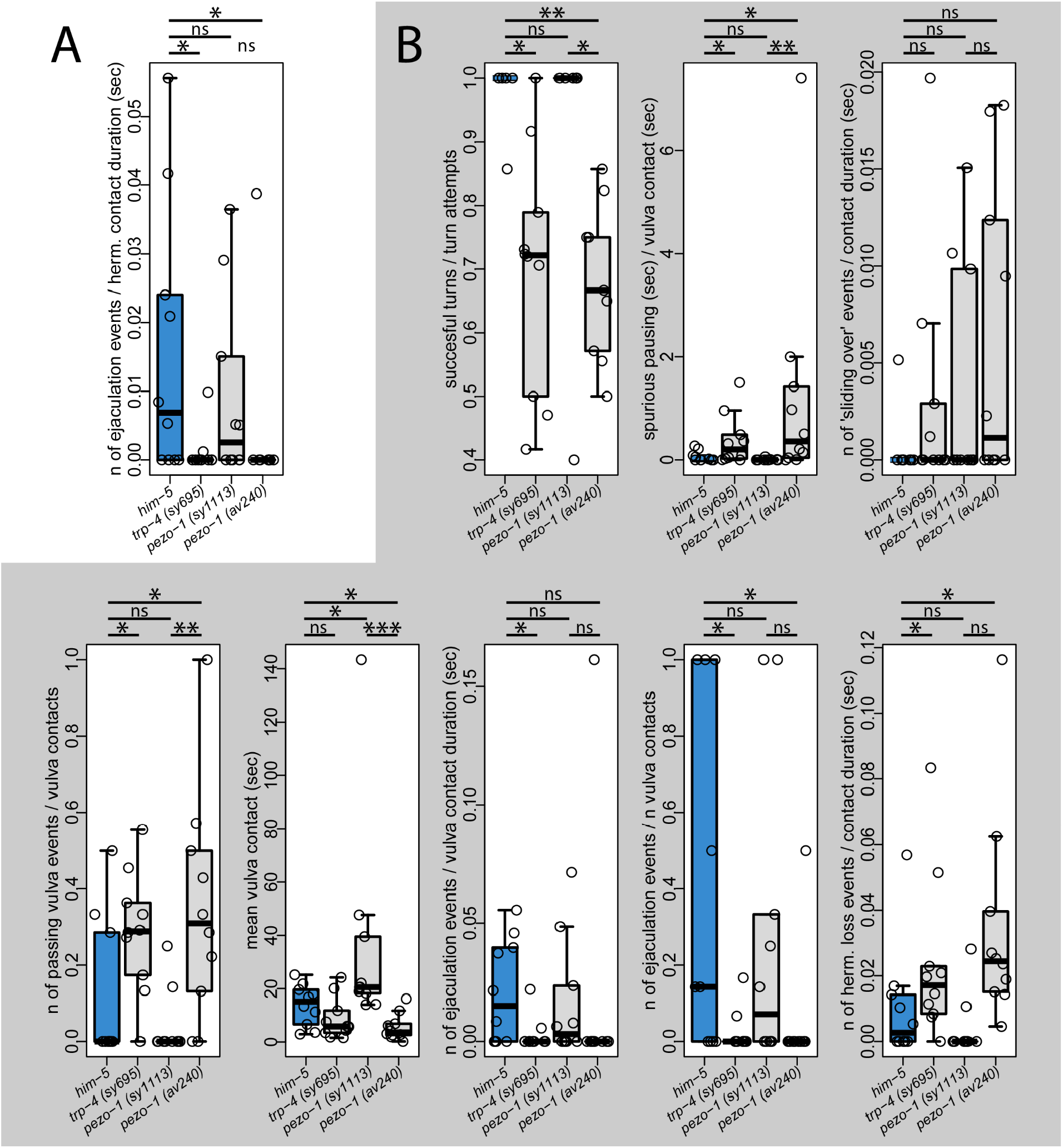
Mating phenotypes of mutants with freely moving hermaphrodites. (A) The *pezo-1(av240)* mutant – a full deletion of the *pezo-1* gene – and *trp-4(sy695)* have low mating efficiency compared to control males *(him-5(e1490))* and C-terminal *pezo-1* deletion, *pezo-1(sy1113)*. (B) Multiple steps of mating are compromised in *pezo-1(av240)* and *trp-4(sy695)* mutants. These mutants have fewer successful turns per turning attempt, more spurious pausing, more vulva passes, and fewer ejaculation events per time spent at the vulva. All mutant strains in this experiment were crossed with *him-5(e1490)*. Ten males of each strain were recorded. Wilcoxon test, ^*^p<0.05, ^**^p<0.01, ^***^p<0.001, ns – not significant.

### Multiple motifs of mating are compromised in *pezo-1* and *trp-4* mutants

Low mating efficiency can result from defects in any one of the many steps that comprise overall mating behavior. To pinpoint the phenotypes of *pezo-1* and *trp-4* mutants, we analyzed video recordings of mating behavior, focusing on specific behavioral motifs: scanning, turning, stopping at the vulva, spurious spicule insertion attempts away from the vulva, sliding over the hermaphrodite’s lateral side, and sperm release. We found that *pezo-1(av240)* and *trp-4(sy695)* mutants, but not *pezo-1(sy1113)* were significantly less efficient at multiple motifs of mating behavior. Compared to control males, *pezo-1(av240)* and *trp-4(sy695)* mutants were less likely to complete successful turns (Wilcoxon test, *p<0*.*011, p<0*.*004* respectively), showed more spurious spicule insertion attempts away from the vulva (Wilcoxon test, *p<0*.*039, p<0*.*027*), and were less likely to stop at the vulva (Wilcoxon test, *p<0*.*043, p<0*.*048*). When *pezo-1(av240)* and *trp-4(sy695)* males did reach the vulva, they were less likely to perform the full spicule insertion and sperm release (Wilcoxon test, *p<0*.*044, p<0*.*026*) (Figure 2B). *pezo-1(av240)* males also lost the hermaphrodite more easily (Wilcoxon test, *p<0*.*007*). Mutations in both genes compromise multiple steps of mating, and result in partially-overlapping phenotypes.

### Full deletion of *pezo-1* has a stronger phenotype than a C-terminal deletion

The *pezo-1(av240)* mutant (a complete deletion of *pezo-1*) had much stronger mating defects compared to the *pezo-1(sy1113)* mutant (a 2kb in-frame C-terminal deletion). The C-terminal deletion in *pezo-1(sy1113)* removes the pore domain of PEZO-1 (Figure 1C). One interpretation is that piezo with the large C-terminal deletion retains some function. Consistent with this hypothesis, a prior study in hermaphrodites showed that the full deletion *pezo-1(av240)* has a much stronger effect on the brood size compared to *pezo-1(sy1199)*, a mutant with a Crispr/Cas-9 knock-in of a “STOP-IN” cassete^62^ that prevents C-terminus translation^26^. We tested if the hermaphrodite brood size is also different between *pezo-1(av240)* and *pezo-1(sy1113)* mutants. Indeed, brood size comparisons show that the *pezo-1(sy1113)* mutants have a significantly larger brood size than the *pezo-1(av240)* mutants, ∼80 progeny per animal vs ∼20 progeny per animal over a three day period (Supplemental Figure 2). Complete deletion of *pezo-1* has a significantly larger effect on both mating efficiency and egg laying compared to a C-terminal deletion, which affects the pore domain.

### *pezo-1* mutant males are more likely to fail turn attempts but show normal PVV activity upon successful turns

*pezo-1* is strongly expressed in the PVV interneuron (Figure 1I-K). Based on the results of functional imaging, ablation phenotypes, and synaptic wiring, PVV has been implicated in turning behavior^43,63^. The male performs turning via sharp ventral bending of his tail when he reaches either end of the hermaphrodite. In the wild type males, most turning attempts are successful and end with the tail attaching itself to the other side of the hermaphrodite, allowing the male to continue scanning for the vulva (Figure 2B). In the *pezo-1 (av240)* mutant, 32% ± 12%(SD) turn attempts fail and the tail loses contact with the hermaphrodite. We tested whether PVV showed different activity upon turning attempts in the wild type males and *pezo-1 (av240)* mutants. PVV activity was extracted and aligned to the onset of turning attempts. Successful turns and failed turns were analyzed separately. In the wild type, PVV activity starts rising shortly before the turn, and peaks 1.2 seconds (median) after the turn onset (Figure 3A). In the *pezo-1 (av240)* mutants, PVV activates upon turning and its temporal dynamics is similar to the wild type (PVV peaks at 1.4 seconds, Figure 3B). In the *pezo-1 (av240)* mutants, more than 50% of turning attempts were unsuccessful during brain-wide imaging experiments. In both wild type and mutant males PVV peaks later – at 1.8 seconds – when the turning is unsuccessful (Figure 3C-E). Although *pezo-1 (av240)* mutants show compromised turning, and compromised turning is associated with a delay in PVV activation (both in mutants and wild type), it is uncertain if the *pezo-1* mutation itself causes the delayed activation of PVV.

**Figure 3.**
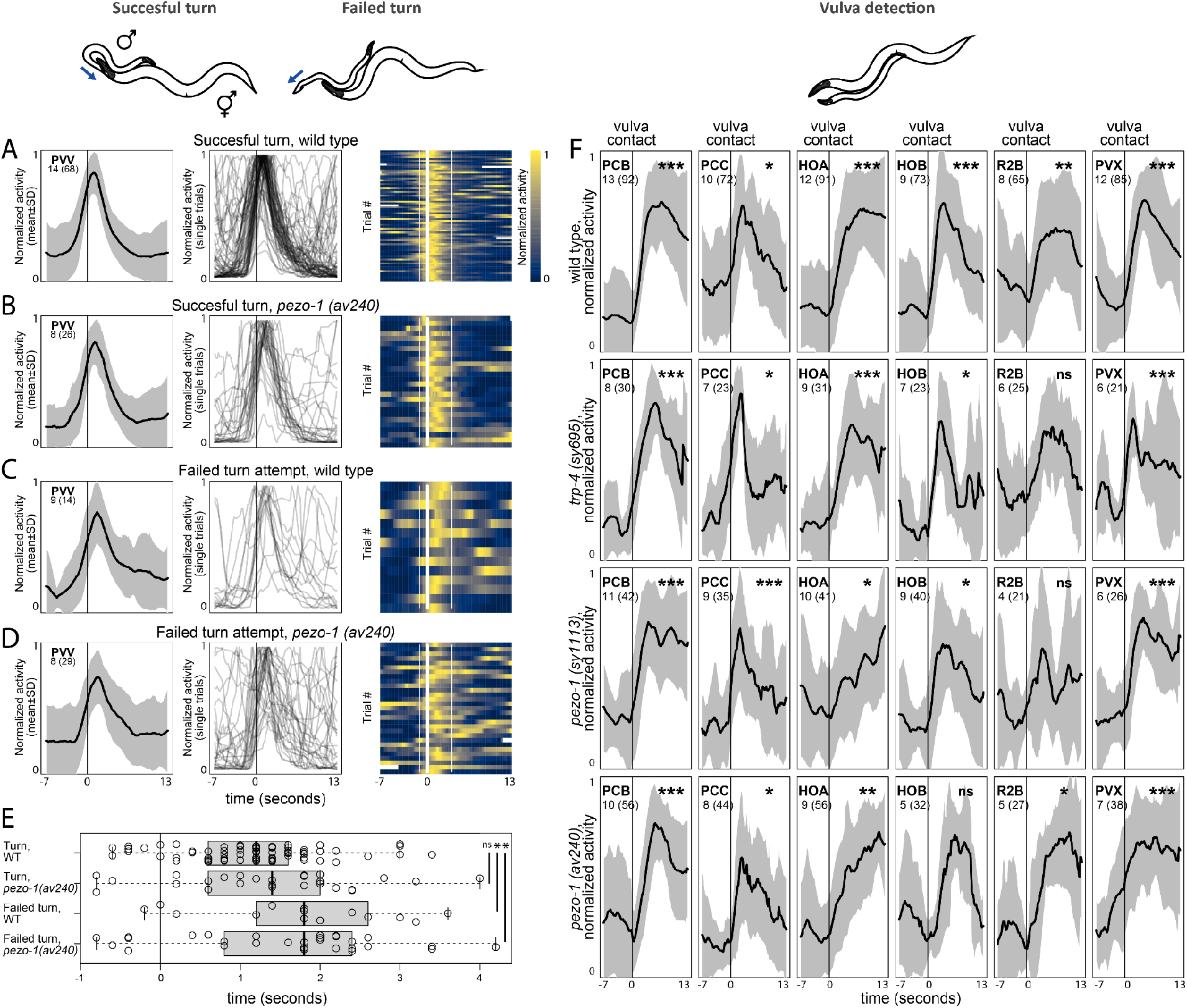
Activities of neurons aligned to behavioral events in *pezo-1* and *trp-4* mutants. (A) PVV activates with the onset of turning. Mean dataset-averaged activity is shown as well as PVV traces for individual turning attempts. (B-D) *pezo-1* mutants show similar PVV dynamics to wild type males during both successful and failed turning attempts. (E) PVV activates later during failed turning attempts in both wild type and *pezo-1* males. Peak activation time relative to the turn onset is shown; Wilcoxon test, ^*^p<0.05, ns – not significant. (F) Activities of neurons of the vulva-detecting circuit aligned to the onset of vulva detection. Vulva detection involves a circuit of recurrently-connected neurons, including two postcloacal sensilla neurons PCB and PCC, the hook neurons HOA and HOB, the R2B ray neuron, and the PVX interneuron. In the wild type control, these neurons become active when the tail reaches the vulva. In *pezo-1(sy1113), pezo-1(av240)*, and *trp-4(sy695)* activation of PCB, PCC, HOA, HOB, R2B, and PVX appears to be largely uncompromised. Black traces show normalized mean dataset-averaged activities across all animals; standard deviation is indicated in gray. The number of animals is shown; the number of events recorded across all animals is shown in parentheses. One-sample t-test, ^*^p<0.05, ^**^p<0.01, ^***^p<0.001, ns – not significant.

### Neurons for vulva detection show normal activity upon vulva detection in *pezo-1* and *trp-4*

Given that *pezo-1* and *trp-4* are expressed in neurons involved in vulva detection, we tested whether neurons in the previously identified vulva-detecting circuit – PCB, PCC, HOA, HOB, R2B, and PVX – exhibited different activity upon vulva contact in mutant males compared to control males. We recorded the activity of the male’s entire posterior brain during mating with freely-moving hermaphrodites. We extracted activities of PCB, PCC, HOA, HOB, R2B, and PVX and aligned them to the onset of vulva-detecting events. In control males, PCB, PCC, HOA, HOB, R2B, and PVX show a sharp increase in activity when the male tail contacts the vulva (Figure 3F). The same neurons show a similar increase in activity upon vulva contact in *pezo-1(sy1113), pezo-1(av240)*, and *trp-4(sy695)* mutants (Figure 3F). Thus, in *pezo-1* and *trp-4* mutants, the response of individual neurons of the vulva-detecting circuit to vulva detection events is largely unaffected.

### Neurons for vulva detection show spurious activation away from the vulva in *pezo-1* and *trp-4*

*pezo-1(av240)* and *trp-4(sy695)* mutants frequently pause scanning away from the vulva and show spurious spicule insertion attempts (Figure 2B). Previously, pausing away from the vulva was shown to be accompanied by pulses of spurious activation in the vulva-detecting neurons^43^. We screened recordings of *pezo-1(sy1113), pezo-1(av240)*, and *trp-4(sy695)* mutants for such spurious activation of the vulva-detecting circuit (Figure 4). In control males, spurious activation is uncommon. Two of twenty-four (9%) control males showed brief pulses of spurious activation. In contrast, spurious activity was observed in 5 of 19 (26%) *pezo-1(sy1113)* males, 5 of 10 (50%) *pezo-1(av240)* males, and 13 of 15 (86%) *trp-4(sy695)* males.

**Figure 4.**
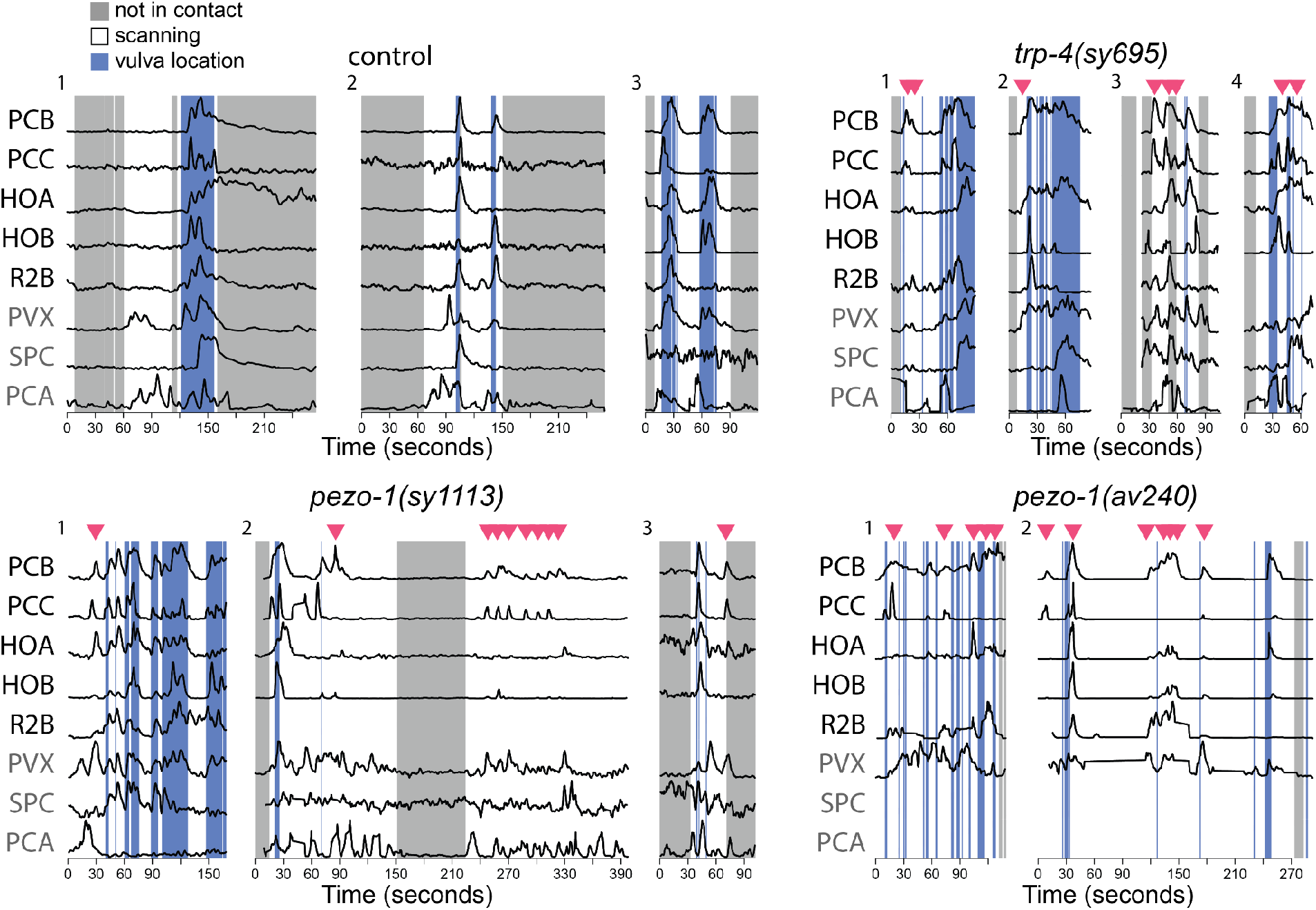
*pezo-1* and *trp-4* mutants show spurious activation among vulva-detecting neurons. Normalized activity traces of neurons involved in vulva detection, copulation, and scanning in wildtype, *trp-4*, and *pezo-1* males. In the control males, sensory neurons PCB, PCC, HOA, HOB, and R2A activate specifically when the male passes over the vulva and are inactive when scanning, 5 of 19 (26%) *pezo-1(sy1113)* males, 5 of 10 (50%) *pezo-1(av240)* males, and 13 of 15 (86%) *trp-4(sy695)* males exhibited spurious activation during scanning. In the figure, examples of spurious activation in the mutant males are shown. Red triangles indicate bursts of spurious activation. Activities of additional neurons are shown in gray: PVX, an interneuron which non-specifically activates with vulva location, SPC, which activates with spicule insertion, and PCA which activates with backward sliding.

### PCA role in mechanosensation is compromised in *trp-4* mutants

Previously it was suggested that PCA mechanosensory activation with backward scanning inhibits spurious activation of the vulva-detecting circuit^43^. When PCA is ablated, the vulva-detecting circuit becomes active away from the vulva^43^. The molecular mechanism of this inhibition is likely glutamatergic^58^. PCA is presynaptic to several neurons involved in vulva detection, including HOA, PCB, and PVX^63,64^. Given that *trp-4* is expressed in the PCA neuron, we asked if the spurious activity in *trp-4* mutants might be caused by the loss of mechanosensory function of PCA and the consequent release of the vulva-detecting circuit from glutamatergic inhibition from PCA.

In control males, the activity of PCA is correlated with the relative sliding between the male and the hermaphrodite. When the male tail slides backwards relative to the hermaphrodite, PCA increases its activity (t-test, p<0.001). When the male tail slides forward – PCA activity decreases (Figure 5A) (t-test, p<0.001). In contrast, in *trp-4* males, PCA activity does not change when the direction of the relative sliding changes (Figure 5A) (t-test, p<0.528 and p<0.342). Also, unlike in control males, in *trp-4* mutants PCA does not become active upon ventral contact with the hermaphrodite (Figure 5B) (t-test, p<0.001 for control and p<0.762 for *trp-4* males). This suggests that mechanosensation in PCA is dependent on the TRP-4 channel. Without functioning TRP-4, PCA cannot monitor the relative sliding between the male and hermaphrodite and is thereby unable to provide glutamatergic inhibition to vulva-detecting neurons during scanning. Lack of inhibition results in the spurious activation of vulva-detecting neurons.

**Figure 5.**
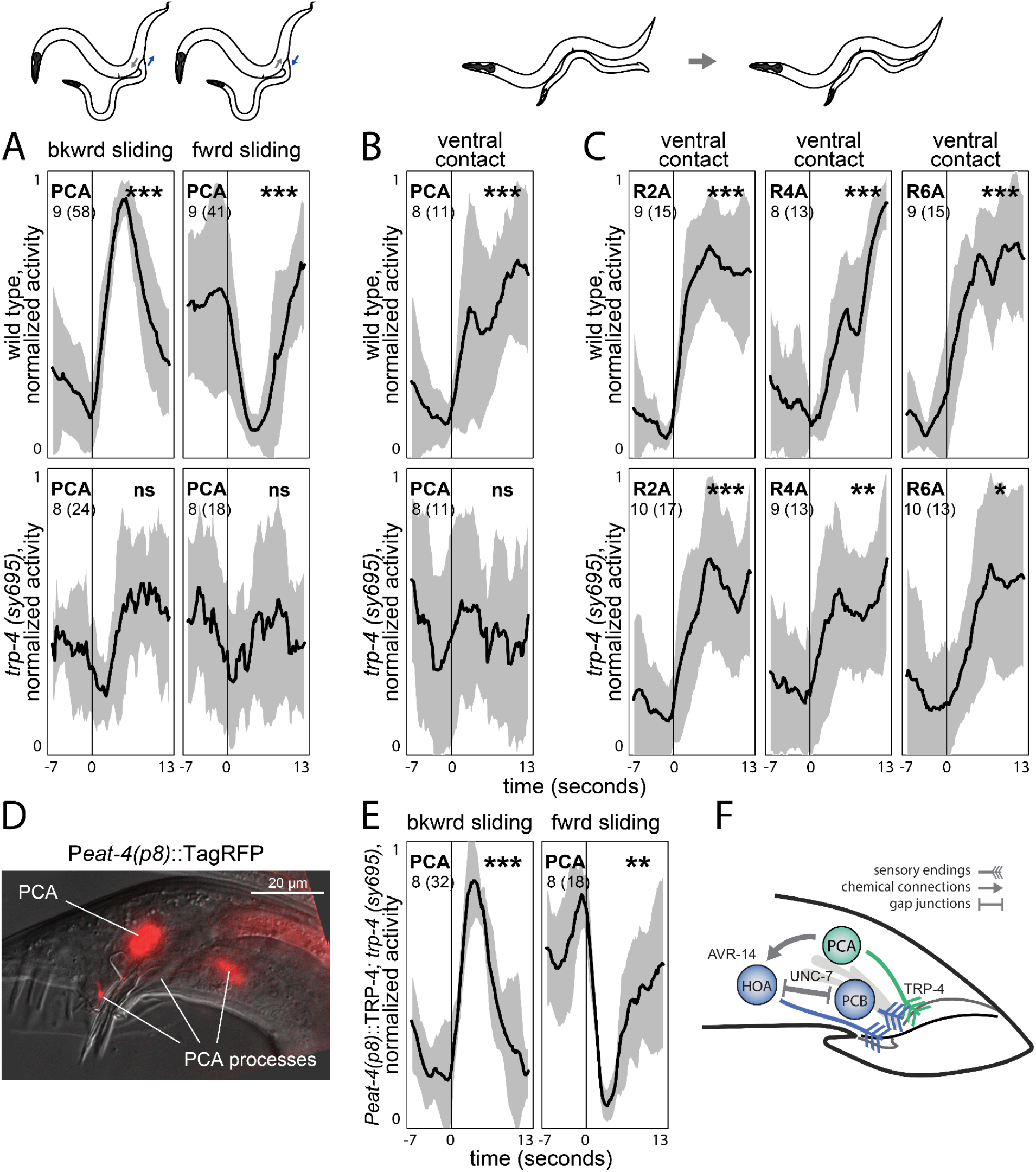
*trp-4* role in mechanosensation. (A) In control males, PCA activity increases when the male tail slides backwards relative to the hermaphrodite, and decreases when the tail slides forwards. In *trp-4* mutants no change in activity occurs. (B). Control males, but not *trp-4* mutants exhibit activation of PCA upon ventral contact with the hermaphrodite. A-type ray neurons show activation upon ventral contact in both control and *trp-4* mutant males (C). (D) A 289 bp promoter 5327 bp upstream of the *eat-4* start codon (*eat-4(p8)*) drives expression in PCA and HOA. Right lateral aspect; PCA and its processes are shown. (E) Cell-specific expression of TRP-4 using *eat-4(p8)* promoter (PS9669) restores PCA sensitivity to sliding. (F) PCA’s mode of action. PCA uses TRP-4 to sense when the male tail slides backwards relative to the hermaphrodite. Glutamatergic PCA, activated with sliding, hyperpolarizes HOA via inhibitory glutamate-gated chloride channel AVR-14. Hyperpolarization of HOA can propagate to vulva-detecting neurons via electrical synapses (UNC-7). TRP-4 mutation compromises PCA’s mechanosensation, thereby releasing vulva-detecting neurons from inhibition and resulting in their spurious activation away from the vulva. Black traces show normalized mean dataset-averaged activities across all animals; standard deviation is indicated in gray. The number of animals is shown; the number of events recorded across all animals is shown in parentheses. One-sample t-test, ^*^p<0.05, ^**^p<0.01, ^***^p<0.001, ns – not significant.

### A-type ray neurons show normal activity in *trp-4* mutants

Like the PCA neuron, A-type ray neurons express *trp-4* and have been implicated in mechanosensation. Many A-type ray neurons increase their activity upon ventral contact with a hermaphrodite. We tested whether the A-type ray neurons in *trp-4* mutants are less activated by the contact. We extracted activities of R2A, and R6A, and aligned them to the onset of the ventral contact between the male and the hermaphrodite. We also extracted activities of R4A, which does not express *trp-4*^59^. Like R2A and R6A, R4A increases its activity upon contact (Figure 5C). In *trp-4* males R2A, R4A, and R6A all showed similar activity, and like in control males, these ray neurons were activated upon contact (Figure 5C), although the activation appeared weaker compared to control males. These results are consistent with previous observations that *trp-4* males show normal hermaphrodite response^59^. We conclude that A-type ray neurons do not rely solely on TRP-4 for hermaphrodite recognition.

### Cell-specific TRP-4 expression restores PCA sensitivity to sliding in *trp-4* mutants

We tested if PCA mechanosensation, compromised in *trp-4* mutants, can be rescued by cell-specific expression of TRP-4. We created a rescue line using a 289 bp region of the *eat-4* promoter^52^, which drives gene expression in two neurons in the tail – PCA and HOA (Figure 5D). We performed brain-wide imaging of the rescue line during mating and quantified PCA activity in response to relative sliding between mating partners. In the rescue line, PCA showed wild type-like responses to changes in the sliding direction (Figure 5E). PCA activity increased when the male slided backwards relative to the hermaphrodite and it decreased when the male switched to sliding forwards. We conclude that expression of TRP-4 in PCA is sufficient to restore PCA’s mechanosensory role in monitoring relative movement between mating partners.

## DISCUSSION

Our results show that both PEZO-1 and TRP-4 play important roles in *C. elegans* mating. Mutations in both genes reduce mating efficiency by altering the performance of multiple motifs of mating. Multiple behavioral phenotypes exhibited by the mutants is consistent with the broad expression of *pezo-1* and *trp-4* throughout male nervous system. Both genes are expressed in neurons that participate in many different motifs of mating. For example, *pezo-1* is strongly expressed in neurons that contribute to two non-overlapping mating motifs: the PVV interneuron, which was previously shown to be required for turning behavior, and the PCB sensory neuron, which is involved in vulva detection.

Behavioral phenotypes of *pezo-1* and *trp-4* partially overlap. This is also consistent with the broad expression patterns of both *pezo-1* and *trp-4*. Even when the two genes are expressed in different neurons, the neurons themselves might contribute to the same behavioral motifs. For example, both PVV (which expresses *pezo-1*) and A-type ray neurons (which express *trp-4*) have been implicated in turning^40,43^.

In the case of PEZO-1 we were unable to fully dissect mechanistic links between mutant alleles and observed behavioral phenotypes. In the case of TRP-4, however, we were able to identify its key contribution to PCA’s ability to monitor sliding between the male and hermaphrodite, which is used to regulate switching between two behavioral motifs – scanning and stopping at the vulva.

Our results suggest that PEZO-1 that has a mutation in its pore domain might retain some of its function. Mutants with complete deletion of *pezo-1* (*pezo-1(av240)*) demonstrate fewer ejaculation events, successful turns, and vulva contacts than a partial deletion, while also causing more spurious pausing, “pass vulva” events, and “loss of contact” events than the *pezo-1(sy1113)* mutants. Consistent with this, a larger deletion of *pezo-1* has a more severe phenotype on a neuronal level: *pezo-1(av240)* males show more events of spurious activation of the circuit involved in vulva detection. In addition, *pezo-1(sy1113)* has a less severe egg laying phenotype compared to *pezo-1(av240)*, a result consistent with Bai et al. 2020^26^, where a less severe egg laying phenotype was reported for the “STOP-IN” allele in the pore domain. These results are surprising because previous electrophysiological studies of mutant piezo channels showed the key role of the pore domain in determining ion-conducting properties^17,65–67^. In a recent *C. elegans* study Hughes et al. (2022)^28^ showed that mutations affecting C-terminus of PEZO-1 had a weaker effect on the defecation frequency but stronger effect on the egg-laying rate compared to mutations at the 5’ end of the gene. We cannot rule out completely that *pezo-1(av240)* phenotypes might be affected by another genome feature in the *pezo-1* locus, which might have been altered by the large deletion. However, given that the non-pore containing region of the human and mouse Piezo1 has been demonstrated previously to be sufficient for mechanotransduction^22,66^, it is plausible that *C. elegans* PEZO-1 with the C-terminal deletion preserves some of its function.

Our results provided further evidence for a mechanosensory role of PCA in male mating behavior and suggested that its function relies on TRP-4 as the key mechanotransducer (Figure 5F). Previous recordings from PCA in freely-moving males during mating revealed that PCA plays distinct roles in copulation^43,51^ and in monitoring the relative sliding between the male and the hermaphrodite^43^. We show that *trp-4* is expressed in PCA. In *trp-4* mutants, PCA does not become active when the male slides backward relative to the hermaphrodite. Moreover, *trp-4* mutants pause backward scanning away from the vulva, which recapitulates the phenotype caused by PCA ablation. Cell-specific TRP-4 expression in the PCA neuron restores its ability to monitor sliding between mating partners. We suggest that initiation of backward sliding of the male tail relative to the hermaphrodite activates the TRP-4 channel, which in turn activates PCA. This activation of PCA provides glutamatergic inhibition to the neurons involved in vulva detection, which is required for persistent scanning^43,58^.

PCA has been implicated in multiple motifs of mating. In addition to the monitoring of backward sliding, PCA has been demonstrated to play roles in copulation^43,51^, and previous ablation studies indicated its contribution to vulva detection^40,61^. Whether these other roles of PCA require TRP-4 remains to be studied.

Many *pezo-1* and *trp-4* mutant phenotypes are incompletely penetrant. For example, PCB, which expresses *pezo-1*, shows nearly normal activity in both *pezo-1* mutant alleles; A-type ray neurons R2A and R6A, which express *trp-4*, retain their activation by the hermaphrodite contact in *trp-4* mutants. The incomplete penetrance of the phenotypes might be due to additional, yet uncharacterized components that may contribute to activation of these neurons in response to mechanical as well as chemical stimuli from the hermaphrodite. Furthermore, many sensory neurons in the posterior brain of *C. elegans* male are connected to each other, a property that is thought to enable recurrent excitation. It is plausible that even though mutant PEZO-1 or TRP-4 might be incapable of activating a neuron in response to a relevant mechanical stimulus from a hermaphrodite, the neuron itself might still be activated by other sensory neurons, which are tuned to correlated inputs from the mating partner.

## ACKNOWLEDGEMENTS

We thank WormBase for genome information, CGC for providing strains, Ian Kimbell for help with data collecting, Xiaofei Bai for providing AG570 *pezo-1(av240)*, Shawn Xu for sharing a plasmid containing *trp-4*, and Tsui-Fen Chou for providing Cas9 protein. This work was supported by NIH R01-NS113119 (PWS and ADTS). KB was supported by NIH-Training Grant T32GM007616.

## SUPPLEMENTAL FIGURES

**Supplemental Figure 1.**
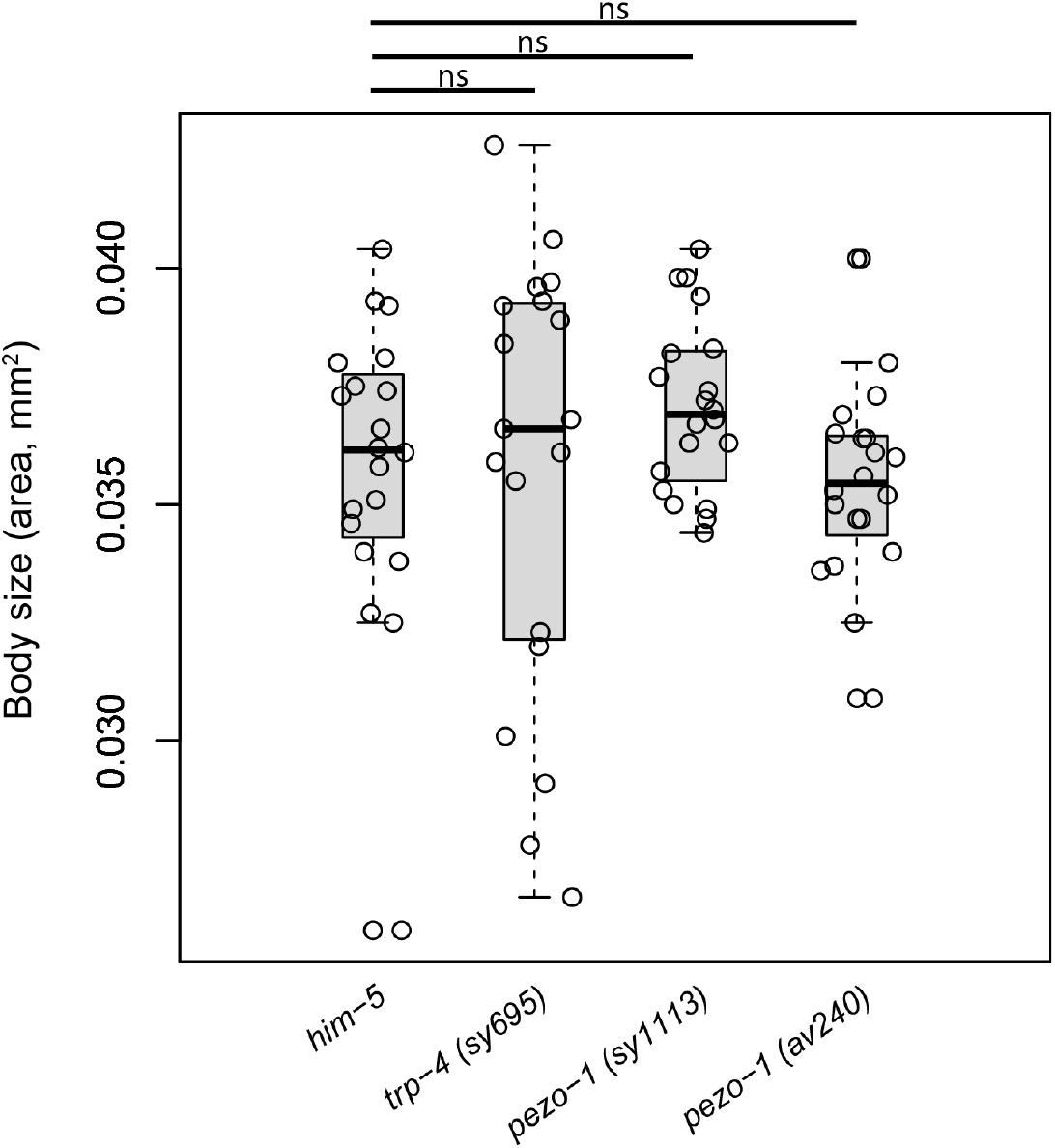
Body size measurements of the wild type and mutant *C. elegans* males. Sagittal area of the wild type and mutant *C. elegans* males. t-test, the significance threshold was set to 0.05.

**Supplemental Figure 2.**
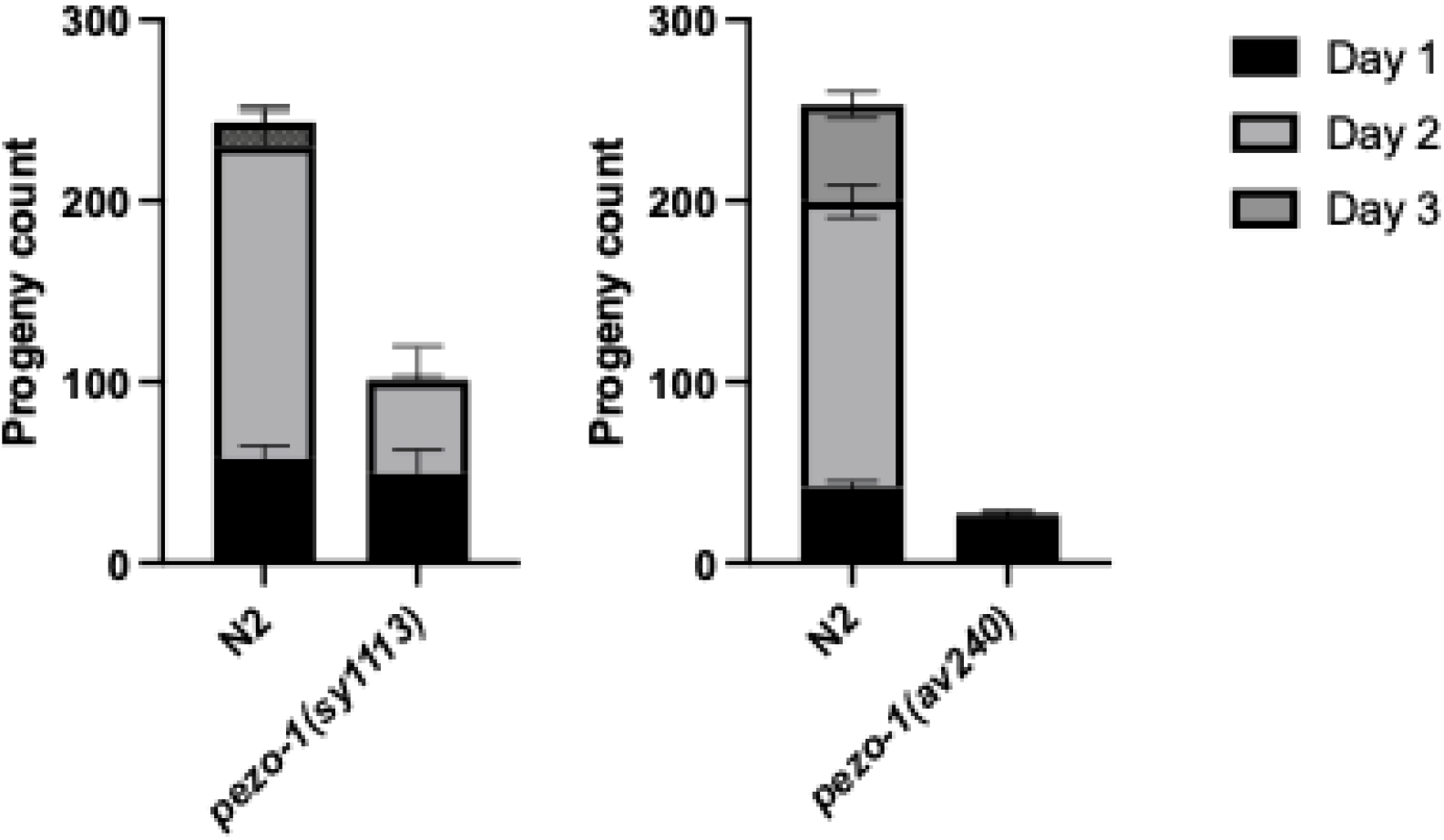
Brood size reduction in *pezo-1(sy1113)* and *pezo-1(av240)* mutants. In control animals, brood size over the course of three days was normal at around 250 progeny spanning that period. For mutant animals, both *pezo-1(sy1113)* and *pezo-1(av240)* exhibited significantly reduced brood sizes. However, the brood size for *pezo-1(sy1113)* was on average around 80 animals while the brood sizes for *pezo-1(av240)* was on average around 20 animals. Thus *pezo-1(av240)* has a much stronger brood size reduction than *pezo-1(sy1113)*. These results are consistent with previous results by Bai et al^26^.

